# U-PASS: unified power analysis and forensics for qualitative traits in genetic association studies

**DOI:** 10.1101/605766

**Authors:** Zheng Gao, Jonathan Terhorst, Cristopher Van Hout, Stilian Stoev

## Abstract

**Summary:** Despite the availability of existing calculators for statistical power analysis in genetic association studies, there has not been a model-invariant and test-independent tool that allows for both planning of prospective studies and systematic review of reported findings. In this work, we develop a web-based application U-PASS (Unified Power analysis of ASsociation Studies), implementing a unified framework for the analysis of common association tests for binary qualitative traits. The application quantifies the shared asymptotic power limits of the common association tests, and visualizes the fundamental statistical trade-off between risk allele frequency (RAF) and odds ratio (OR). The application also addresses the applicability of asymptotics-based power calculations in finite samples, and provides guidelines for single-SNP-based association tests. In addition to designing prospective studies, U-PASS enables researchers to retrospectively assess the statistical validity of previously reported associations.

**Availability and implementation:** U-PASS is available as a web-based R Shiny application at https://power.stat.lsa.umich.edu. Source code is available at https://github.com/Pill-GZ/U-PASS.

**Supplementary information:** Supplementary data are available in the application.

## 1. Introduction

The probability of detecting a true association between genetic and phenotype variations, known as statistical power, is influenced by a number of factors such as sample sizes, the statistical test used, the frequency of the risk variant, and magnitude of the effect on the trait. Power analysis, which determines the suitable factor combinations in order to achieve sufficient statistical power, plays an important role in determining study designs (Skol *et al*., 2003; Goodwin *et al*., 2005), and in interpreting published findings (Ioannidis, 2005).

There has been a number of widely used calculators for genome-wide association studies (GWAS). Early work by Sham (1998) studied power analysis of likelihood ratio tests for associations between marker SNPs and quantitative or qualitative traits; the results were implemented in GPC (Purcell *et al*., 2003). Skol *et al*. (2003) studied the performance of two-sample t-tests, and extended the analysis to two-stage designs; the results were implemented in the CaTS caclulator, and later, in the GAS calculator for one-stage designs (Johnson *et al*., 2017). Independently, Menashe *et al*. (2008) implemented the calculations for one-stage designs in the PGA calculator. Recent works have also studied power of a number of SNP-set based tests targeting rare variants (Wang *et al*., 2014; Derkach *et al*., 2017). See Sham & Purcell (2014) for a review.

Despite these efforts, some difficulties remain in practice:

### 1. Lack of universality

Existing power analyses are tied to the underlying models and the statistical procedures used; power calculations for a certain model-method combination may not be valid if either the model or the method changes. In principle, power calculations based on likelihood ratio tests or t-tests cease to hold for studies running logistic regressions or chi-squared tests. Users are burdened with matching the appropriate tool to the specific type of analysis they wish to perform. This is complicated by the fact that the precise test and model assumptions are rarely made explicit in the existing calculators.

### 2. Mismatching definitions of key quantities

While GWAS catalogs, e.g., NHGRI-EBI (MacArthur *et al*., 2016), require studies to report risk allele frequency (RAF) *in the control group*, all of the aforementioned power calculators assume the RAF input as the frequency *in the general population*. These quantities are not necessarily equal, and using one in place of the other may grossly distort power estimates.

### 3. (In)accuracies in finite samples

While existing tools rely on large-sample approximations in their power calculations, these approximations are not reliable in finite samples when genetic variants are rare. Existing calculators are silent about the applicability of asymptotics-based approximations, and how they should be corrected.

As a result, it is not only challenging to use the existing power calculation tools for planning genetic association studies correctly, but also difficult to systematically review the statistical validity of findings reported in the literature, since different models and tests must be handled differently, and with care.

In an effort to address these difficulties and deficiencies, we propose a unified framework for power analysis of single variant association studies. By abstracting away the assumptions of disease models and testing procedures which may vary from study to study, we reduce the problem to the essential quantities that are invariant to nuisance parameters. These ideas are implemented in the software U-PASS, enabling model-invariant, test-independent power analysis, as well as systematic reviews of the statistical validity of reported findings.

We briefly summarize the important features and uses of the software below. Mathematical details and results from numerical experiments are collected in the Supplement.

## 2. Methods and features

### 2.1 A model-invariant parametrization

We identify the core parameters common to models of qualitative traits,

- Sample sizes, i.e., the number of Cases, *n*_1_, and Controls, *n*_2_,
- Conditional distribution of risk variant among Controls, i.e., risk allele frequency (RAF) in the Control group, denoted *f*.
- Odds ratio (OR) of genetic variants, denoted R.

An alternative hypothesis, e.g., a disease model, determines the set of quantities either implicitly or explicitly, and therefore determines statistical power for a given test procedure. Here, statistical power is calculated directly based the quartet (*n*_1_, *n*_2_, *f, R*), allowing us to perform power analysis valid for studies employing different models.

We make the important distinction between RAF in the Control group (*f*), versus RAF among all subjects in the study, and RAF in the general population. Throughout this work and the software, RAF refers to the risk allele frequency in the Control group, consistent with the reporting standard of the NHGRI-EBI Catalog.

### 2.2 A test-independent power analysis

While power calculations are necessarily tied to the statistical tests used, for many common association tests, statistical powers are asymptotically equivalent. We show in the Supplement that the likelihood ratio test, chi-square test, Welch’s t-test, and LR test for logistic regressions have asymptotically the same power, as long as the core parameters assume the same values. Our model-invariant, test-independent analysis allows us to calculate power in a unified fashion. The formulas used for power calculations in terms of the core parameters are detailed in Section 1 of the Supplementary material.

Our software only requires users to specify the number of cases and controls. The common power limits are calculated as a function of RAF and OR, given numerically, and visualized as a heatmap in the OR-RAF diagram; see Figure 1 for a preview of the software graphical user interface.

**Fig. 1.**
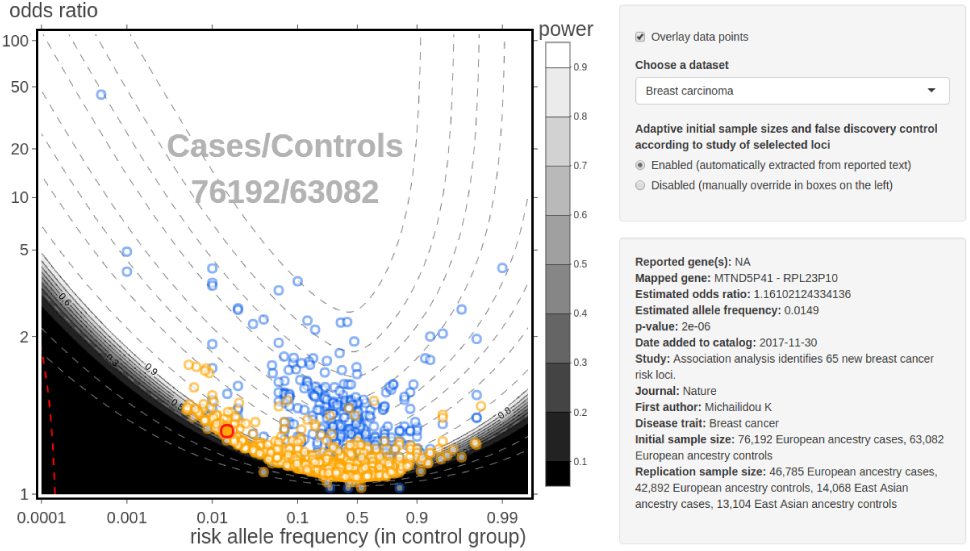
Snapshot of the application’s user interface, displaying reported associations from breast cancer studies in the NHGRI-EBI GWAS Catalog (circles). The findings are overlaid on the OR-RAF power diagram of association tests (greyscale heatmap). The initial sample sizes are dynamically adjusted, and automatically determined from texts of the article reporting the user selected loci (red circle). Information of the selected loci and the articles are also dynamically displayed; findings reported in the same article are highlighted (orange circles). We provide finite sample corrections by marking the rare variant region(s) where asymptotic approximations do not apply (red dashed lines, lower-left). The majority of the published findings we surveyed exhibited a striking level of concordance with our theoretical predictions, with most associations congregating just inside the detectable region.

### 2.3 Review and forensics of reported findings

This unified treatment allows us to examine results from different studies in the same diagram, even when they do not employ the same model or statistical test. This enables systematic reviews of reported findings for their statistical validity. In particular, a reported association predicted to have low power given the study’s sample size – lying in the dark regions of the OR-RAF diagram – while not impossible, invites further scrutiny.^1^

Studies where reported associations show misalignment with the predicted powered curves may be further investigated for potential problems in the data curation process. We reached out to one study where gross misalignment was identified (Dominguez-Cruz *et al*., 2014). The authors of the study confirmed that this was due to a problem in the data curation process when the findings were added to the GWAS Catalog (Dominguez Cruz, personal communication).

The software provides options for users to load and overlay findings reported in the NHGRI-EBI GWAS Catalog, or upload data from other sources compliant with the Catalog’s data format.

### 2.4 Rare variants and finite sample corrections

We address the quality of asymptotic approximations in our power analysis, as well as the applicability of single variant tests when rare genetic variants are present. Specifically, we provide a lower bound on the variant counts needed to calibrate Fisher’s exact test. If variant counts fall below the threshold, exact tests, and by extension, single-SNP-based association tests, cannot be correctly calibrated to have the desired type I error rate without sacrificing all statistical power. In such cases, the asymptotic approximations do not apply. We mark this low-count, low-power region on the OR-RAF diagram. We refer readers to Section 2 in the Supplement for further details.

The software also provides options for users to specify the rare variant as 1) having less than a specified counts, or 2) occurring in less than a percentage of all subjects in the study, as is customary in the literature.

## 3 Implementation

U-PASS is implemented as an interactive web-based R Shiny application, hosted at https://power.stat.lsa.umich.edu, open to the public. It has been designed to be user-friendly and self-contained. Source code is freely available at https://github.com/Pill-GZ/U-PASS.

## Footnotes

1 It should be noted that a reported association predicted to have high power is not automatically accurate, as winner’s curse induced by multiple testing may inflate the OR and RAF estimates (Xiao & Boehnke, 2009).

