## Supplement for "U-PASS: unified power analysis and forensics for qualitative traits in genetic association studies"

**Abstract**

We provide details of the unified analysis of statistical power for common association tests in Section 1. We also elaborate on a simple rule for finite-sample correction of the asymptotic calculations in Section 2. The performance of this correction is examined through numerical experiments in Section 4. Proof of the main result is provided in Section 3.

### 1 Unified asymptotic analysis of power

In association tests for categorical phenotypes and genetic variants, counts of subjects in each phenotype-genetic variant combination are tabulated in the form of a contingency table. For a 2-genetic-variant-by-2-phenotype definition, we have the following table of counts.

| | Genotype | | |
|---|---|---|---|
| # Observations | Variant 1 | Variant 2 | Total by phenotype |
| Cases | $O_{11}$ | $O_{12}$ | $n_1$ |
| Controls | $O_{21}$ | $O_{22}$ | $n_2$ |

Statistics are then calculated based on the counts, to test for associations between the genotypes and phenotypes, at levels adjusted for multiplicity. Performance of a test is measured in terms of power, i.e., probability of correct rejection under an alternative hypothesis.

Power analysis starts by assuming an alternative, typically described by a disease model (recessive, dominant, additive, etc.). Power of a test is approximated either based on large sample asymptotics, or through simulating the empirical distribution of the statistic under the alternative.

Note that even when the disease model dictates more than two genotypes (e.g., one heterozygous and two homozygous variants in an additive model), the association tests may still be based on only two derived variants. This can be due to either grouping of the genotypes definitions in dominant or recessive models, or by adopting a direct comparison of the proportions of allele types between the Case and Control groups, as opposed to the proportions of zygosity. Indeed, the latter approach is the basis of power calculations in Skol *et al.* (2003).

#### 1.1 A model-invariant parametrization

Regardless of the disease model assumed, the test statistics are calculated based on cell counts of the contingency table. The alternative hypothesis, consequently, influences the distribution of the test statistics only through altering the distribution of the multinomial integer counts in the contingency tables.

Consider 2-by-2 multinomial distributions with probability matrix $\mu = (\mu_{ij})_{2\times 2}$,

| | Genotype | | |
|---|---|---|---|
| Probabilities | Variant 1 | Variant 2 | Total by phenotype |
| Cases | $\mu_{11}$ | $\mu_{12}$ | $\phi = \mu_{11} + \mu_{12}$ |
| Controls | $\mu_{21}$ | $\mu_{22}$ | $1 - \phi = \mu_{21} + \mu_{22}$ |

We may assume – by relabelling, and hence without loss of generality – that genetic Variant 1 is positively associated with the Cases, and referred to as the risk allele/variant.

The multinomial probability matrix $\mu$ can be fully parametrized by the parameter triple:

- Fraction of Cases $\phi$, i.e., marginal distribution of phenotypes.
- Conditional distribution of risk variant among Controls, i.e., risk allele frequency (RAF) in the Control group
$$f := \mu_{21}/(1-\phi). \tag{1}$$
- Odds ratio (OR) of the genotype Variant 1 to Variant 2
$$\mathrm{R} := \frac{\mu_{11}}{\mu_{21}} \Big/ \frac{\mu_{12}}{\mu_{22}} = \frac{\mu_{11}\mu_{22}}{\mu_{12}\mu_{21}}. \tag{2}$$

An alternative hypothesis, e.g., a disease model, determines the the trio of quantities either implicitly or explicitly, and therefore fully determines statistical power for a specific test at a given sample size. From a statistical perspective, the disease model serves no role except to specify the distribution of the counts under the alternative. Power may be calculated by directly prescribing the trio $(\phi, f, R)$, and the sample size.

It is worth pointing out that while disease models play no role beyond specifying the alternative, they do sometimes inform our choice of a test statistic, hence influencing statistical power in higher order contingency tables. These tests include, e.g., the Cochran-Armitage test, and variations thereof; see González *et al.* (2008); Li *et al.* (2008) for further examples where tests are tailored to disease models.

We make the important distinction between RAF *in the Control group* ($f$), versus RAF *in the study* ($\mu_{11} + \mu_{21}$), and RAF *in the general population.* Throughout this work, RAF refers to the risk allele frequency in the Control group, consistent with the reporting standards of the NHGRI-EBI Catalog (MacArthur *et al.*, 2016).

#### 1.2 Conditional vs unconditional tests

Readers familiar with the underlying assumptions of association tests in contingency tables may have noticed that we have described a multinomial distribution of the cell counts. That is, we have only conditioned on the total number of observations in the study. This is indeed the assumption behind tests such as (the original, unconditional version of) the likelihood ratio test, and the Person chi-square test.

It is not, however, the assumption behind some other tests. For example, analysis of t-tests typically assumes the observed number in each arm of the study are given. That is, we would condition on the phenotype marginals when comparing the proportions of genetic variants among the Cases and Controls. It is perhaps an assumption most close to reality, where the number of samples collected in each arm of the study are pre-determined. In this case, we have two binomial observations, Binom$(n_1, p_1)$ and Binom$(n_2, p_2)$, instead of a multinomial observation.

| | Genotype | | |
|---|---|---|---|
| Cond. Prob. | Variant 1 | Variant 2 | Counts by phenotype |
| Cases | $p_1$ | $1 - p_1$ | $n_1$ |
| Controls | $p_2$ | $1 - p_2$ | $n_2$ |

RAF and OR can be similarly defined,

- Marginal distribution of Cases, fixed at $\phi := n_1/(n_1 + n_2)$,
- Risk allele frequency (RAF) in the Control group is the synonymous with the conditional distribution of risk variant among Controls,

$$f := p_2,$$

- Odds ratio (OR)

$$\mathrm{R} := \frac{p_1\phi}{p_2(1-\phi)} \Big/ \frac{(1-p_1)\phi}{(1-p_2)(1-\phi)} = \frac{p_1(1-p_2)}{(1-p_1)p_2}.$$

Alternative hypotheses may be formed as in the multinomial case with parameters $\phi$, $f$ and $R$, along with the total samples size.

Finally, we mention the assumptions behind the Fisher's exact test. The Fisher exact test conditions on the number of observations of both the phenotype variants and genetic variants, leading to a hypergeometric distribution of the first cell count $O_{11}$ given the marginals $n_1$, $n_2$, and $O_{11} + O_{21}$, under the null hypothesis. We found no easy parametrizations of alternative hypotheses under this framework. Indeed, existing power calculations for Fisher's exact test resort to simulations under the the two-binomial assumptions (Smyth *et al.*, 2017).

We refer interested readers to the recent work by Ripamonti *et al.* (2017) and Choi *et al.* (2017) which elucidate the controversies regarding the choices of conditioning when performing statistical inferences on 2-by-2 contingency tables. We do not attempt to resolve the controversies in this work. Our goal is to state clearly the assumptions behind the tests, and and show the asymptotic equivalence in terms of power, under their respective assumptions.

#### 1.3 A test-independent power analysis

We now present the main result allowing a unified power analysis, applicable for a wide range of common association tests in 2-by-2 tables.

If we consider a fixed parameter values of $(f, R)$ under the alternative, no matter how close to the null, the probability of rejection of the null hypothesis by any reasonable test should approach one as sample size increases ($n \to \infty$). On the other hand, the probability of rejection is less than one in finite samples, making this type of asymptotics useless for approximation.

Therefore, in order to find finer approximations of power, we study alternatives close to the null. In particular, we take a sequence of alternatives approaching a limit point in the null space, in the hope that limiting rejection probability is between 0 and 1. It turns out – see, e.g., Ferguson (2017) Chapter 10, and Lehmann (2004) Chapter 5 – that the appropriate rate at which the alternatives should shrink towards the limit point is $1/\sqrt{n}$.

Under the multinomial assumption, let $\mu^{(0)}$ be the probability matrix of the (independent) 2-by-2 multinomial distribution, with marginals $(\theta, 1-\theta)$ for the genetic variants, and marginals $(\phi, 1-\phi)$ for the phenotypes. We require that $\theta \in (0,1)$ and $\phi \in (0,1)$ be bounded away from 0 and 1. Let $\mu = \mu^{(n)}$ be the sequence of alternatives such that

$$\sqrt{n}(\mu^{(n)} - \mu^{(0)}) \to \delta \begin{pmatrix} 1 & -1 \\ -1 & 1 \end{pmatrix}, \tag{3}$$

where $\delta$ is a positive constant.

Equivalently, under the two-binomial assumption, let $p_1^{(0)} = p_2^{(0)} = \theta$ be the null hypothesis, with fixed marginals $(\phi, 1-\phi)$ for the phenotypes. Let $(p_1, p_2) = p^{(n)}$ be the sequence of alternatives such that

$$\sqrt{n}\phi(p_1 - \theta) \to \delta \quad \text{and} \quad \sqrt{n}(1-\phi)(p_2 - \theta) \to -\delta, \tag{4}$$

where $\delta$ is a positive constant. It is easy to see that with the same $\delta$ the two sequences of alternatives have the same RAF and OR, and therefore have the same expected number of observations in each cell.

**Theorem 1** *In 2-by-2 contingency tables, under the assumption that the counts in the contingency table follow the multinomial distributions.*

- *The likelihood ratio test for independence,*
- *the likelihood ratio test for zero slope in logistic regressions,*
- *Person's chi-squared test for Independence,*

*and under the assumption that counts in the contingency table follow the two binomial distributions,*

- *the two-sided Welch's t-test for equal proportions*

*have the same asymptotic power curves. Specifically, for the sequence of alternatives defined in* (3) *and* (4)*, all of the listed tests, at level* $\alpha$*, have statistical powers converging to*

$$\mathbb{P}[\chi^2(\lambda) \geq q_\alpha], \tag{5}$$

*where* $q_\alpha$ *is the upper* $\alpha$ *quantile of the central chi-square distribution, and* $\chi^2(\lambda)$ *is a non-central chi-square distribution with non-centrality parameter*

$$\lambda = \delta^2/(\theta(1-\theta)\phi(1-\phi)). \tag{6}$$

The proof of Theorem 1 is detailed in Section 3 below.

Theorem 1 is the central result that paves the way for a unified powers analysis. It allows us to chart findings from different studies employing the applicable tests in the same diagram, with the same power limits. In particular, for large samples, tests for zero slopes in logistic regressions should report approximately the same set of loci as Welch's t-tests for equal proportions on the same dataset, after the same family-wise error rate adjustments. The estimated odds ratios (in the case of logistic regression, estimate slopes exponentiated) and RAF's, when charted on the OR-RAF diagram, should also follow the same power limits.

To use this result for power calculations, we start with an alternative hypothesis, defined by the core parameters $(\phi, f, R)$, and sample size $n$.

| | Genotype | |
|---|---|---|
| Probabilities | Variant 1 | Variant 2 |
| Cases | $fR\phi/(fR+1-f)$ | $(1-f)\phi/(fR+1-f)$ |
| Controls | $f(1-\phi)$ | $(1-f)(1-\phi)$ |

Elementary algebra yields $\theta = fR\phi/(fR+1-f) + f(1-\phi)$, and $\delta = \sqrt{n}(\theta - f)(1-\phi)$. If tests are based on allele type counts, accounting for the fact that each genetic location has a pair of alleles, the effective sample sizes should be doubled, and the appropriate non-centrality parameter becomes $\delta = \sqrt{2n}(\theta - f)(1-\phi)$. Power may then be approximated using the formula in (5).

#### 2 Finite-sample corrections

The result of our power calculations above are only accurate to the extend that the asymptotic approximations are applicable. In practice, of course, we have only finite samples, and the asymptotic approximations no longer hold when cell counts are low. While existing tools have completely ignored this issue, we offer here a simple correction in finite samples by resorting to exact tests.

Specifically, we calculate the minimum number of observations of the genetic variants needed for Fisher's exact test to be correctly calibrated, referred to as the **minimum calibration numbers**. As we shall see in simulations in Section 4, they provide a useful lower bound on the variant counts necessary for any asymptotic approximations to apply.

For a contingency table with marginal phenotype counts $(n_1, n_2)$, and marginal genetic variant counts $(m_1, m_2)$, we calculate the p-values of the most extreme observations according to Fisher's exact test. For rare risk alleles, this corresponds to the following table.

| | Genotype | | |
|---|---|---|---|
| # Observations | Variant 1 | Variant 2 | Counts by phenotype |
| Cases | $m_1$ | $n_1 - m_1$ | $n_1$ |
| Controls | 0 | $n_2$ | $n_2$ |

If the p-values do not fall below the desired type I error threshold, then the rejection region (for $O_{11}$) must lie beyond $m_1$. Under the fixed marginal assumptions of Fisher's exact test, no contingency tables with the given marginals can be rejected at the specified level. In other words, we have given up all power to achieve proper type I error control. Therefore, the minimum counts needed for the risk allele count must exceed $m_1$, in order for association tests to have any power.

For rare non-risk alleles, the most extreme observation corresponds to the following table.

| | Genotype | | |
|---|---|---|---|
| # Observations | Variant 1 | Variant 2 | Counts by phenotype |
| Cases | $n_1$ | 0 | $n_1$ |
| Controls | $n_2 - m_2$ | $m_2$ | $n_2$ |

We can similarly determine the minimum number of non-risk allele counts needed to achieve non-zero power at the given type I error target, for a given phenotype marginals $(n_1, n_2)$.

For correctly calibrated tests, an alternative hypothesis with expected variant counts less than the minimum calibration numbers should have power close to zero; asymptotic power approximations do not apply for these alternatives. We correct the asymptotic approximations laid out in Section 1.3 by setting the predicted statistical power for alternatives in this "rare-variant zone" to zero.

We conduct an extensive simulation study to examine the quality of this finite-sample correction in Section 4. We find this simple rule produces accurate corrections, matching up well to the simulated powers of exact tests. In general, the correction kicks in only for small sample sizes.

In the web-based application U-PASS, we mark the "rare-variant zone" with red dashed lines in the OR-RAF diagram. We also provide options for users to specify the rare-variant threshold by absolute number of counts, or as a fraction of the total number of subjects. We find these two options ad-hoc; the minimum calibration numbers approach provides a more theoretically grounded approximation in power calculations for rare-variants.

### 3 Proof of Theorem 1

#### 3.1 Asymptotic equivalence of likelihood ratio tests and the chi-square test

The asymptotic equivalence of the likelihood ratio (LR) test and the chi-square test in 2-by-2 tables can be found in standard texts on asymptotic theory. See, e.g., Ferguson (2017) Chapter 10 and Chapter 24, and Lehmann (2004) Chapter 5; see also, Hunter (2002) for an accessible derivation of the formula (6).

Recall the likelihood ratio statistic in LR test

$$LR = \frac{\sup_{\mu \in H_1} L(\mu)}{\sup_{\mu \in H_0} L(\mu)}.$$

To see the asymptotic equivalence with the LR statistic under logistic regressions with binary predictors, we reparametrize the likelihood as

$$\begin{aligned} L(\mu) &= \mu_{11}^{O_{11}} \mu_{12}^{O_{12}} \mu_{21}^{O_{21}} (1 - \mu_{11} - \mu_{12} - \mu_{21})^{O_{22}} \\ &= \phi^{n_1} (1-\phi)^{n_2} p_1^{O_{11}} (1-p_1)^{O_{12}} p_2^{O_{21}} (1-p_2)^{O_{22}} \end{aligned}$$

where we have omitted the multinomial coefficient. In the latter parametrization, it is easy to show that the maximizers are $\widehat{\phi} = n_1/n$, $\widehat{p_1} = O_{11}/n_1$, and $\widehat{p_1} = O_{21}/n_2$ under the alternative, and $\widehat{\phi} = n_1/n$, $\widehat{p_1} = \widehat{p_1} = (O_{11} + O_{21})/n$ under the null. Therefore, the terms involving $\phi$ cancels in the LR statistic, and the LR statistic coincides with the logistic regressions likelihood ratio, where $p_1$ and $p_2$ are further reparametrized as

$$\begin{aligned} p_1 &= \exp(\beta_0 + \beta_1)/(1 + \exp(\beta_0 + \beta_1)), \\ p_2 &= \exp(\beta_0)/(1 + \exp \beta_0). \end{aligned}$$

Hence, the logistic regressions likelihood ratio follows the same distribution as in the original likelihood ratio test. Notice that this is not an immediate consequence of the invariance property of the likelihood ratio tests. Rather, it follows because $n_1$, $n_2$ are ancillary for inference of the odds ratios.

##### 3.2 Asymptotic equivalence with Welch's t-test

We now work with the two-binomial assumption, conditioning on the phenotype marginals $n_1$, $n_2$, and show that Welch's t-test has asymptotically the same power. Recall the Welch t-statistic

$$t = \frac{\widehat{p_1} - \widehat{p_2}}{\sqrt{\frac{\widehat{p_1}(1-\widehat{p_1})}{n_1} + \frac{\widehat{p_2}(1-\widehat{p_2})}{n_2}}}, \tag{7}$$

where $\widehat{p_1} = O_{11}/n_1$ and $\widehat{p_2} = O_{21}/n_2$. By the (Lindeberg-Feller) central limit theorem, for the sequence of alternatives defined in (4) we have

$$\sqrt{n_i}(\widehat{p_i} - p_i)\big/\sqrt{p_i(1-p_i)} \Rightarrow \mathrm{N}(0,1), \quad \text{for } i = 1, 2.$$

and therefore, by independence of the two binomial distributions, we have

$$t - (p_1 - p_2)\Big/\sqrt{\frac{p_1(1-p_1)}{n_1} + \frac{p_2(1-p_2)}{n_2}} \Rightarrow \mathrm{N}(0,1). \tag{8}$$

By the definition of the alternatives in (4), we know that

$$\sqrt{n}(p_1 - p_2) \to \delta/(\phi(1-\phi)). \tag{9}$$

On the other hand, for the denominator, we have

$$\begin{aligned}
&\sqrt{n}\sqrt{\frac{p_1(1-p_1)}{n_1} + \frac{p_2(1-p_2)}{n_2}} \sim \sqrt{\frac{p_1(1-p_1)}{\phi} + \frac{p_2(1-p_2)}{1-\phi}} \\
&= \left(\frac{(\theta + O(1/\sqrt{n}))(1-\theta+O(1/\sqrt{n}))}{\phi} + \frac{(\theta + O(1/\sqrt{n}))(1-\theta+O(1/\sqrt{n}))}{1-\phi}\right)^{1/2} \\
&\sim \sqrt{(\theta(1-\theta))/(\phi(1-\phi))}.
\end{aligned} \tag{10}$$

Dividing (9) by (10), in view of (8), we conclude that the centers of the distribution of $t$ converges to

$$\delta/\sqrt{\theta(1-\theta)\phi(1-\phi)},$$

which is precisely the square root of the non-centrality in (6). Finally, the conclusion in

Theorem 1 follows from the fact that the square of a normal distribution with mean $\sqrt{\lambda}$ is equal in distribution to a chi-square distribution with non-centrality parameter $\lambda$.

### 4 Numerical illustrations

We examine the accuracy of the asymptotic approximations in Theorem 1 in finite samples, and of the correction by minimum calibration number introduced in Section 2, via numerical simulation.

#### 4.1 Simulation settings

We compare the theoretical results with powers of Fisher's exact test obtained by simulations. We choose to compare against exact tests for their superior performance in finite samples – while approximate tests like the chi-square test and likelihood ratio tests may fail to protect against type I error inflation when sample sizes are small, exact tests maintain the correct levels.

We simulate from the two-binomial model at a range of parameter values

- Sample size total, $n$: $10^2$, $10^3$, $10^4$, $10^5$,
- Fraction of Cases in the study, $\phi$: 5%, 15%, 50%, 85%,
- Risk allele frequencies in control group, $f$: a grid of 100 values ranging from 0.01% to 99.5%,
- Odds ratio, $R$: a grid of 100 values ranging from 1 to 100.
- p-value cutoffs: $5 \times 10^{-5}$, $5 \times 10^{-8}$.

Each of the $4 \times 4 \times 100 \times 100 \times 2$ parameter value combinations were simulated 1000 times. The results for p-value cutoff at $5 \times 10^{-8}$ are visualized with heatmaps, organized by increasing fraction of Cases in the study $\phi$, in Figures 1 to 4.

Left panels of the figures show the theoretical predictions obtained by Theorem 1; right panels show the estimated powers from simulation. Upper panels are for sample sizes 100; middle panels are for sample sizes $1,000$; and lower panels are for sample sizes $10,000$. "Rare-variant zones" are marked in red dashed lines according to the minimum calibration numbers for both theoretical predictions and simulated results.

#### 4.2 Accuracy of asymptotic predictions and finite sample corrections

We comment on some major features of the theoretical predictions next.

We find that the asymptotic approximations in (5) accurate for balanced designs (Figure 3) and designs with over-sampled Cases (Figure 4), even at sample sizes as low as 100. The asymptotic approximations are also accurate for designs with under-sampled Cases when the sample sizes are above $10,000$ (Figure 1 and 2, lower panels).

In small samples, the asymptotic approximations in (5) tend to produce optimistic predictions for designs with under-sampled Cases. This is seen in the upper panels of Figures 1 and 2. Some combinations of low risk allele frequencies and large odds ratios are predicted to be detected with high powers inside the low-count zones according to the asymptotics (left panels, white regions intersect red dashed lines), while simulations show the set of "high-power" alternatives is in fact slightly smaller, and require the expected variant counts to exceed the minimum calibration numbers (right panels, white regions do not intersect the low-count zones).

Fortunately, the mismatch between the asymptotics-based predictions and reality only takes place at very small sample sizes, and only for very large odds ratios. The proposed correction based on minimum calibration numbers seems to provide a simple resolution for this mismatch. Alternatives with expected allele counts less than required by the minimum calibration numbers (outside the red dashed lines) should be interpreted to have near zero power. That is, in the OR-RAF diagrams, regions outside the red dashed lines should also be colored dark. The corrected predictions are closer to what is obtained for exact tests.

For small odds ratios, numerical experiments suggest that our power calculations are robust across a wide range of Case-Control ratios and sample sizes.

Theoretical predictions for p-value cutoff at $5 \times 10^{-5}$ are more accurate than for p-value cutoff at $5 \times 10^{-8}$, but qualitatively similar. For sample sizes at $100,000$, we find the simulated power curves are almost indistinguishable to theoretical predictions. In the interest of space, we only illustrated the more challenging cases.

#### Funding

This work is partially supported by NSF Grant DMS-1830293, Algorithms for Threat Detection.

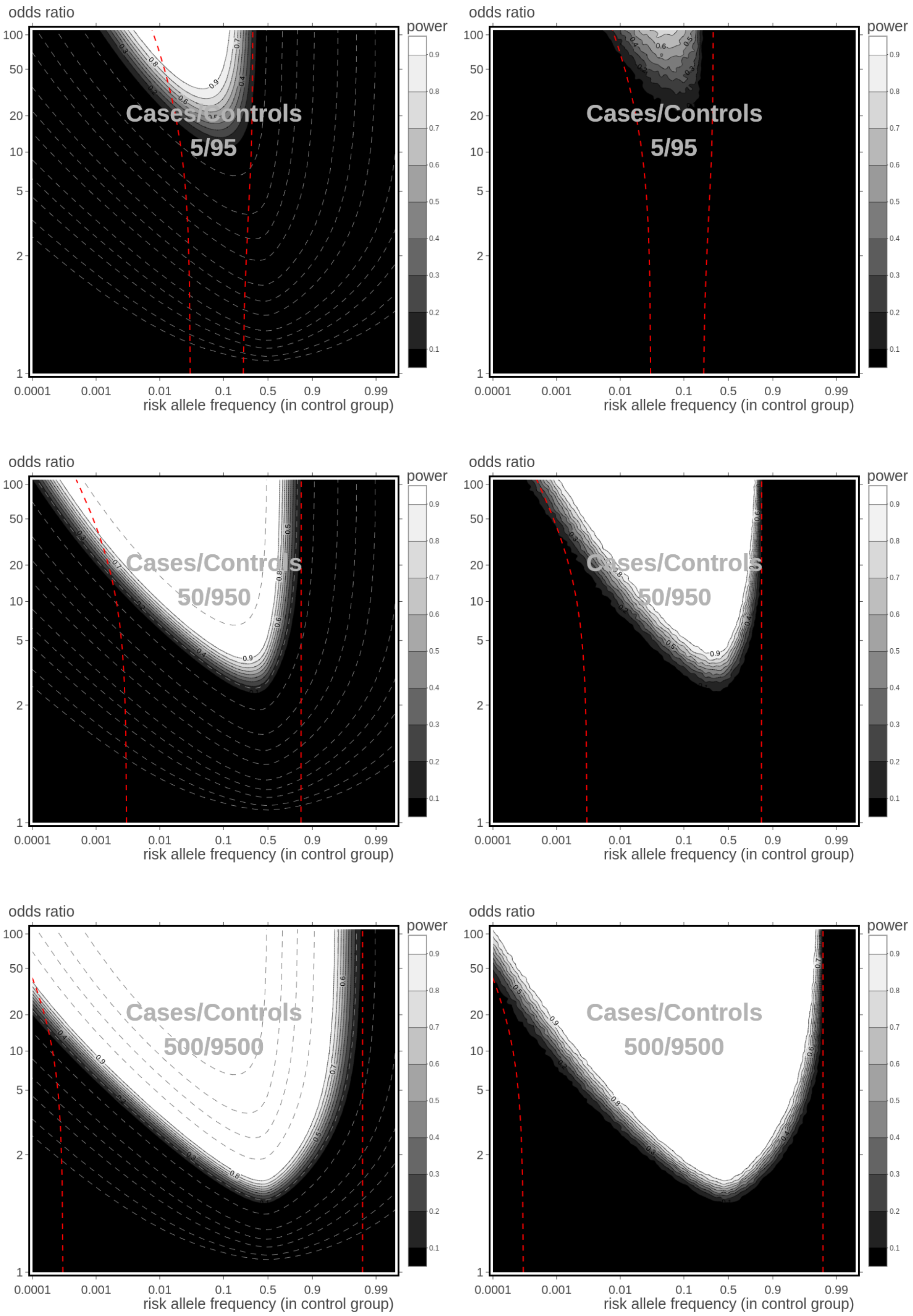

Figure 1: Statistical powers for the OR-RAF combinations, obtained from theoretical predictions (left panels) and by simulations (right panels), for sample sizes $n = 100$ (upper), $n = 1,000$ (middle), and $10,000$ (lower). p-value threshold is at $5 \times 10^{-8}$. Fractions of Cases $\phi = 5\%$.

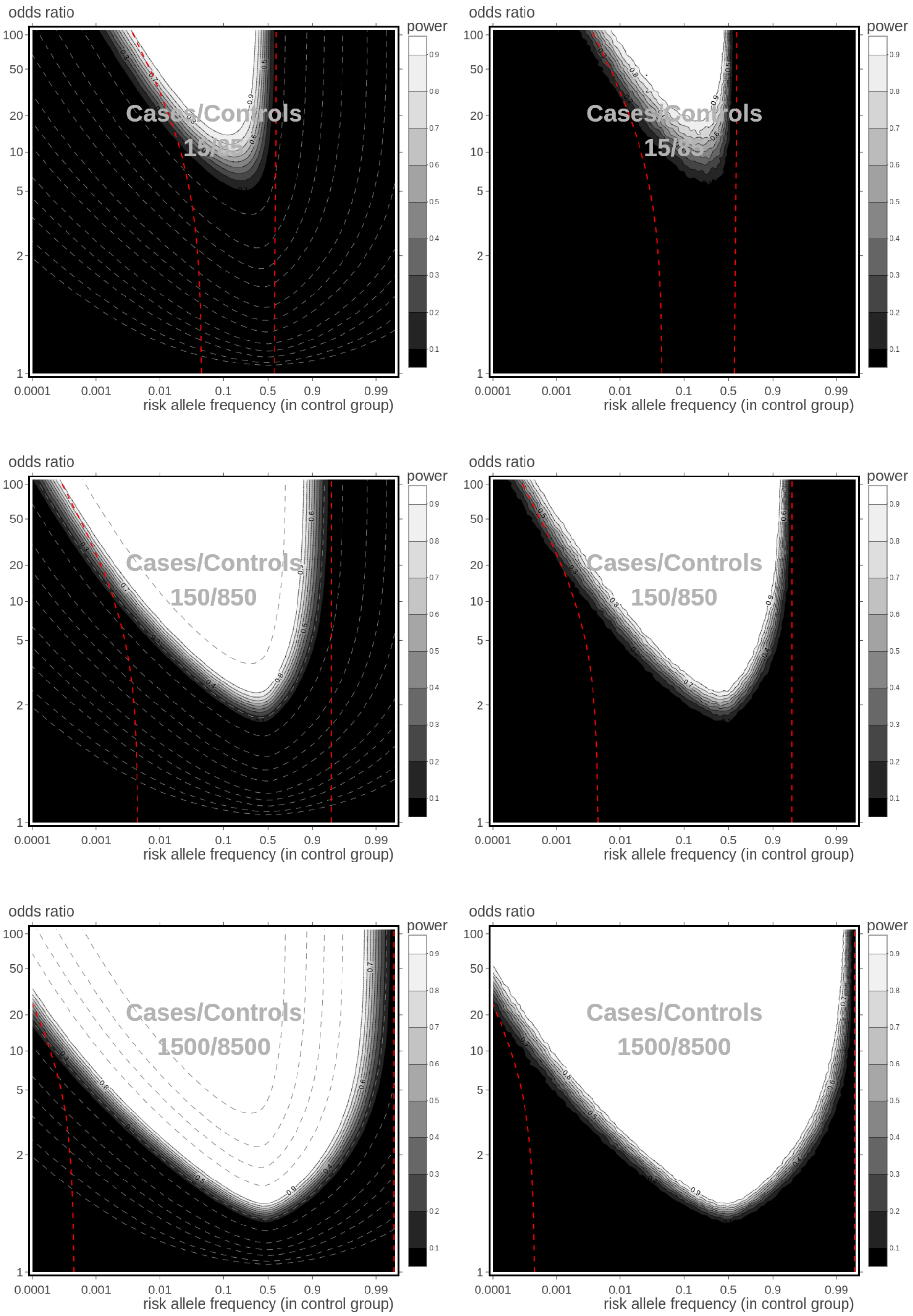

Figure 2: Statistical powers for the OR-RAF combinations, obtained from theoretical predictions (left panels) and by simulations (right panels), for sample sizes $n = 100$ (upper), $n = 1,000$ (middle), and $10,000$ (lower). p-value threshold is at $5 \times 10^{-8}$. Fractions of Cases $\phi = 15\%$.

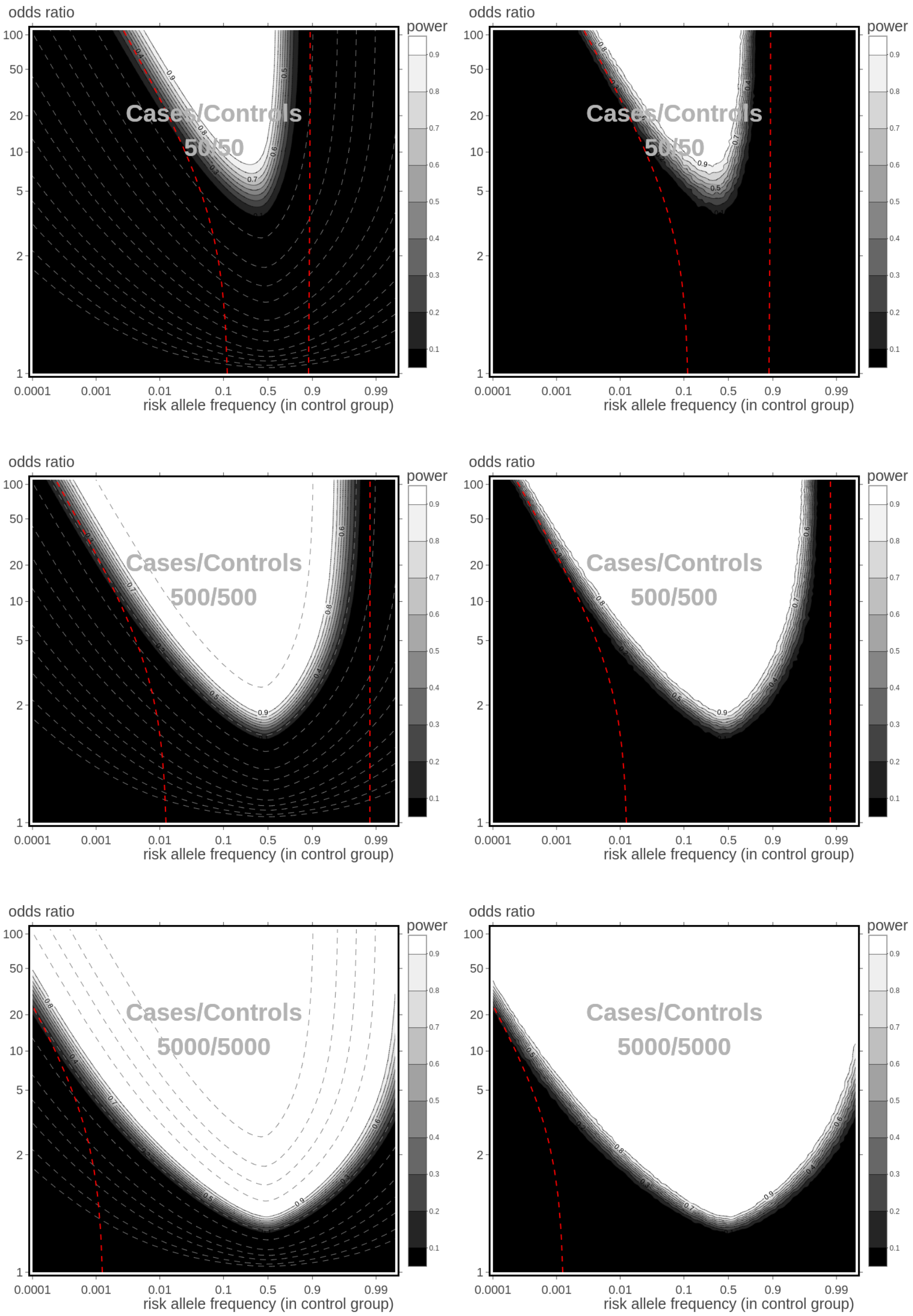

Figure 3: Statistical powers for the OR-RAF combinations, obtained from theoretical predictions (left panels) and by simulations (right panels), for sample sizes $n = 100$ (upper), $n = 1,000$ (middle), and $10,000$ (lower). p-value threshold is at $5 \times 10^{-8}$. Fractions of Cases $\phi = 50\%$.

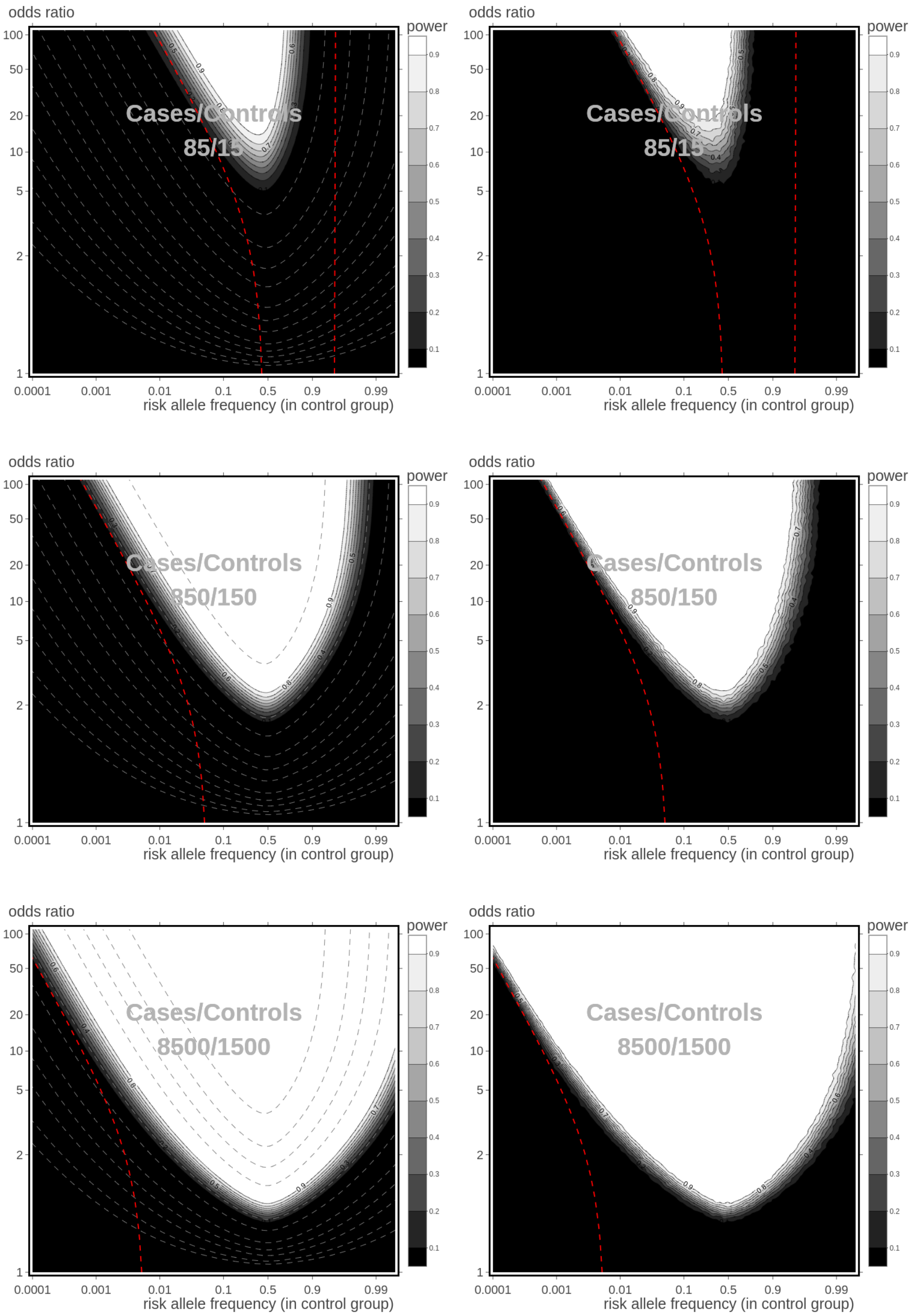

Figure 4: Statistical powers for the OR-RAF combinations, obtained from theoretical predictions (left panels) and by simulations (right panels), for sample sizes $n = 100$ (upper), $n = 1,000$ (middle), and $10,000$ (lower). p-value threshold is at $5 \times 10^{-8}$. Fractions of Cases $\phi = 85\%$.

González, J. R., Carrasco, J. L., Dudbridge, F., Armengol, L., Estivill, X., & Moreno, V. (2008). Maximizing association statistics over genetic models. *Genetic Epidemiology*, **32(3)**, 246-254.

Hunter, D. (2002) Topic 25: Asymptotic Power of Pearsons Chi-Square Test in *Statistics 597A: Asymptotic Tools (Fall 2002), PSU Lecture notes,* http://personal.psu.edu/drh20/asymp/fall2002/lectures/ln11.pdf

Johnson, J. L., & Abecasis, G. R. (2017). GAS Power Calculator: web-based power calculator for genetic association studies. *bioRxiv*, 164343.

Lehmann, E. L. (2004). *Elements of large-sample theory.* Springer Science & Business Media.

Li, Q., Zheng, G., Li, Z., & Yu, K. (2008). Efficient approximation of Pvalue of the maximum of correlated tests, with applications to genomewide association studies. *Annals of human genetics*, **72(3)**, 397-406.

MacArthur, J., Bowler, E., Cerezo, M., Gil, L., Hall, P., Hastings, E., ... & Pendlington, Z. M. (2016). The new NHGRI-EBI Catalog of published genome-wide association studies (GWAS Catalog). *Nucleic acids research*, **45(D1)**, D896-D901.

Menashe, I., Rosenberg, P. S., & Chen, B. E. (2008). PGA: power calculator for case-control genetic association analyses. *BMC genetics*, **9(1)**, 36.

Purcell, S., Cherny, S. S., & Sham, P. C. (2003). Genetic Power Calculator: design of linkage and association genetic mapping studies of complex traits. *Bioinformatics*, **19(1)**, 149-150.

Ripamonti, E., Lloyd, C., & Quatto, P. (2017). Contemporary Frequentist Views of the 2 × 2 Binomial Trial. *Statistical Science*, **32(4)**, 600-615.

Sham, P. C. (1998). *Statistics in human genetics.* Wiley.

Skol, A. D., Scott, L. J., Abecasis, G. R., & Boehnke, M. (2006). Joint analysis is more efficient than replication-based analysis for two-stage genome-wide association studies. *Nature genetics*, **38(2)**, 209.

Smyth, G., Hu, Y., Dunn, P., Phipson, B., & Chen, Y. (2017) statmod: Statistical Modeling. R package version 1.4.30.

Wang, G. T., Li, B., Lyn Santos-Cortez, R. P., Peng, B., & Leal, S. M. (2014). Power analysis and sample size estimation for sequence-based association studies. *Bioinformatics*, **30(16)**, 2377-2378.
